# B-RANKO: An ncRNA Foundation Model with Bidirectional Decoder-Only Architecture

**DOI:** 10.64898/2026.11.12.766109

**Authors:** Filip Vučenik, Rafael Josip Penić, Mile Šikić

## Abstract

Ribonucleic acid (RNA) plays central roles in gene expression, regulation and catalysis, and has emerged as an important target for therapeutic design. While recent RNA foundation models have achieved strong performance on representation learning tasks, relatively few approaches support both native sequence generation and transferable representation learning within a single framework. We present **B-RANKO**, a 150M-parameter bidirectional decoder-only ncRNA foundation model that introduces a novel bidirectional autoregressive attention mechanism, enabling a single architecture to support both *de novo* sequence generation and representation learning. Pretrained on a large corpus derived from RNAcentral, Rfam and GARNET, B-RANKO generates biologically plausible and thermodynamically realistic RNA sequences that exhibit strong novelty relative to their closest matches in the pretraining corpus. We further show that B-RANKO can be efficiently fine-tuned for aptamer generation and achieves competitive performance on BEACON, with particularly strong results on RNA engineering tasks. B-RANKO is publicly available on GitHub at filipvucenik/B-RANKO, together with usage examples and instructions for downloading model weights and datasets.

## 1 Introduction

Ribonucleic acid (RNA) occupies a crucial role in molecular biology through the central dogma, mediating the flow of genetic information from DNA to proteins [1]. However, RNA is far more than a passive intermediary. It can catalyze biochemical reactions [2], regulate gene expression through multiple classes of non-coding RNAs and serve as a programmable molecular substrate in biotechnology and medicine [3–5]. This functional versatility is reflected in diverse applications and mechanisms, ranging from guide RNAs in CRISPR-Cas systems [6] and aptamers for molecular recognition [7] to mRNA-based therapeutics and vaccines [8, 9]. Owing to its synthesizability, programmability and broad functional repertoire, RNA has emerged as an attractive target for computational modeling and engineering.

Despite this promise, some of the clearest recent successes in biomolecular machine learning have come in the field of protein and DNA research. AlphaFold2 [10] showed that deep learning can predict highly accurate protein structures from amino-acid sequences. Similar progress has been made for DNA, where large-scale pretrained models such as Evo 2 [11] support both predictive and generative sequence modeling. By comparison, RNA modeling remains less unified, with representation learning and generation mostly addressed by different models.

Most RNA foundation models to date have been encoder-based and optimized for transferable representation learning. RNA-FM [12] provided an early example of how self-supervised pretraining can produce useful representations for RNA structure and function prediction. Later models, including RiNALMo [13] and AIDO.RNA [14], extended this line of work through model and data scaling and broader evaluation, achieving state-of-the-art performance across multiple RNA structure and function benchmarks.

Generative RNA models differ mainly in how they formulate generation. GenerRNA [15] is a decoder-only model that generates RNA autoregressively. However, its causal attention restricts each position to preceding context, which limits its usefulness for representation learning. EvoFlow-RNA [16] supports both generation and representation learning, formulating non-coding RNA generation as masked discrete diffusion. RNAGenesis [17] combines a pretrained transformer encoder with a Q-former and a causal decoder for task-specific generation. In these models, strong representations are paired with encoder-guided or denoising-based generation, and autoregressive generation with causal context. Whether a single decoder-only model can generate RNA autoregressively and also learn representations that transfer to RNA engineering tasks remains an open question.

To study this question, we introduce **B-RANKO**, a 150M-parameter bidirectional decoder-only ncRNA foundation model, illustrated in Figure 1, that combines a novel bidirectional autoregressive attention mechanism with unified support for de novo sequence generation and transferable representation learning. Our main contributions are as follows:

**Figure 1:**
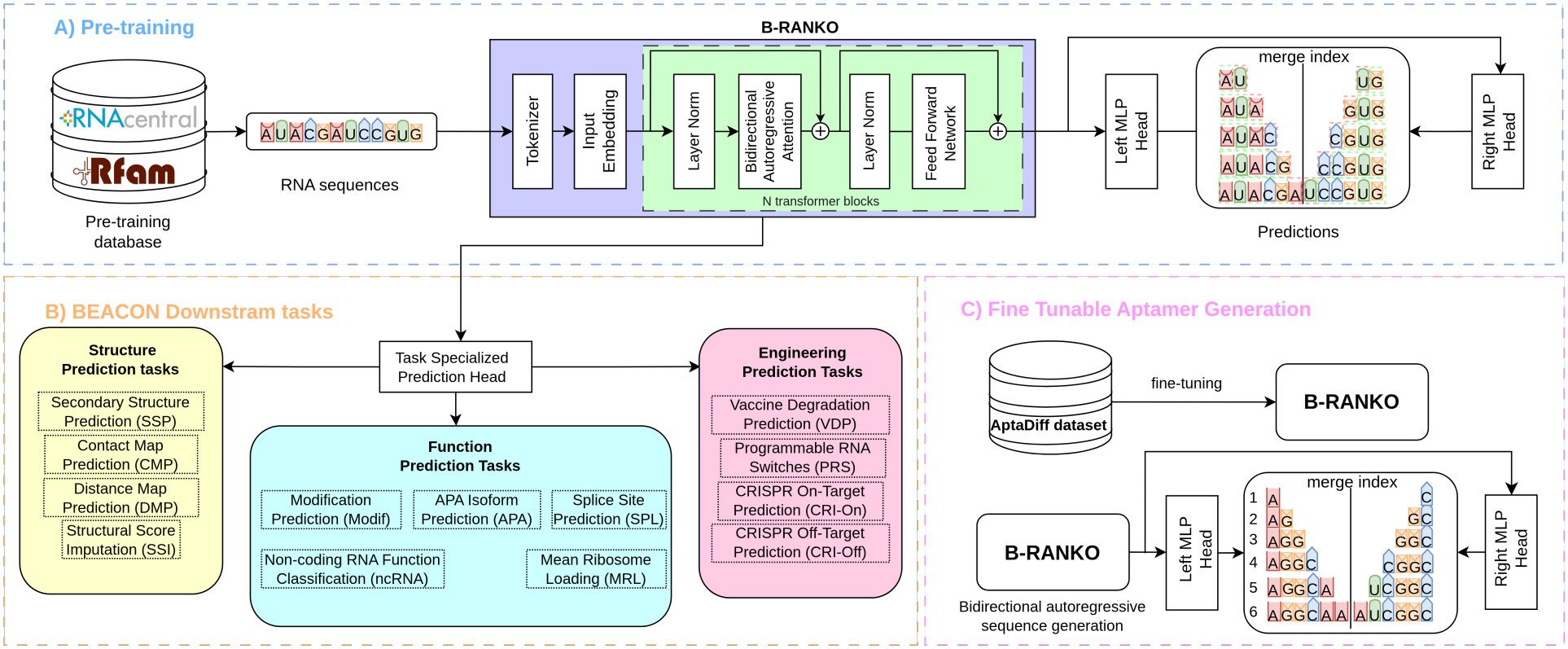
General framework of B-RANKO. A) Pre-training pipeline, model architecture, and training objective. B) Evaluation on BEACON [19] downstream tasks C) Fine-tuning B-RANKO on AptaDiff for bidirectional autoregressive aptamer generation.

- We propose **B-RANKO**, a 150M-parameter bidirectional decoder-only ncRNA foundation model that unifies native sequence generation and transferable representation learning.
- We introduce a **novel bidirectional autoregressive attention mechanism** that enables decoder-only training over both forward and reverse sequence directions, supporting generation and embedding extraction within a single architecture.
- We show that sampling from the pretrained model produces **novel RNA sequences**, assessed by mapping them back to the pretraining corpus with **MMseqs2** [18].
- We demonstrate that B-RANKO can be **fine-tuned efficiently** to generate **aptamer** sequences.
- We evaluate B-RANKO on **BEACON** [19], where it is strongest on **RNA engineering tasks**, with the lowest vaccine degradation error among the compared models.

## 2 Methodology

### 2.1 Model Overview

B-RANKO was developed as the next step after RANKO, our decoder-only autoregressive RNA language model. RANKO was created by adapting the RiNALMo [13] architectural design to a generative setting and replacing bidirectional masked attention with causal attention. This formulation enabled *de novo* RNA generation, but at the cost of losing the bidirectional contextual representations that make encoders effective for downstream prediction tasks. B-RANKO was designed to bridge this gap by introducing a bidirectional decoder-only architecture built around a novel bidirectional autoregressive attention mechanism.

In the experiments that follow, we use RANKO (650M) as a baseline. Its implementation details are given in Supplementary A.1. Because RANKO also differs from B-RANKO in tokenization and size, we isolate the effect of bidirectional attention at matched scale in Section 3.4.

### 2.2 Bidirectional Autoregressive Attention

B-RANKO generates a sequence from both ends toward a meeting point. This design is motivated by RNA secondary structure. Base pairs often join nucleotides that are far apart in the sequence, and the 5^*′*^ and 3^*′*^ ends of RNA molecules tend to lie close together in the secondary structure [20]. In many structured RNAs, such as tRNAs, the two ends pair directly. A left-to-right model generates the 5^*′*^ side of such pairs before any of the 3^*′*^ side exists, whereas inward decoding extends both ends in parallel, so each prediction is conditioned on context from both ends of the sequence.

The idea of generating a sequence in a non-standard order has been explored previously. XLNet uses permutation-based training over different factorization orders [21], while meet-in-the-middle pre-training combines two unidirectional models, with each prediction conditioned on only one side of the sequence [22]. BAnG similarly performs two-front generation for RNA, growing sequences outward from an anchored motif [23]. In contrast, B-RANKO uses a single attention mask parameterized by the merge index *m*. At each decoding step, every prediction can attend to both the generated prefix and generated suffix.

Let an RNA sequence of length *T* be indexed by positions 0, …, *T −* 1. The merge index *m* partitions the sequence into a left prediction region [0, *m*) and a right prediction region [*m, T*). For each query position *i*, B-RANKO defines an inward decoding step:

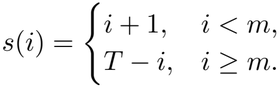

The bidirectional autoregressive mask exposes the prefix generated from the left and the suffix generated from the right at that step:

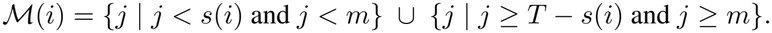

Thus, each query position used in generation attends only to tokens that would already be available under the inward decoding order. As illustrated in Figure 2, this produces an hourglass visibility pattern: positions near the sequence ends observe limited context, while positions closer to the meeting point receive progressively broader context from both sides.

**Figure 2:**
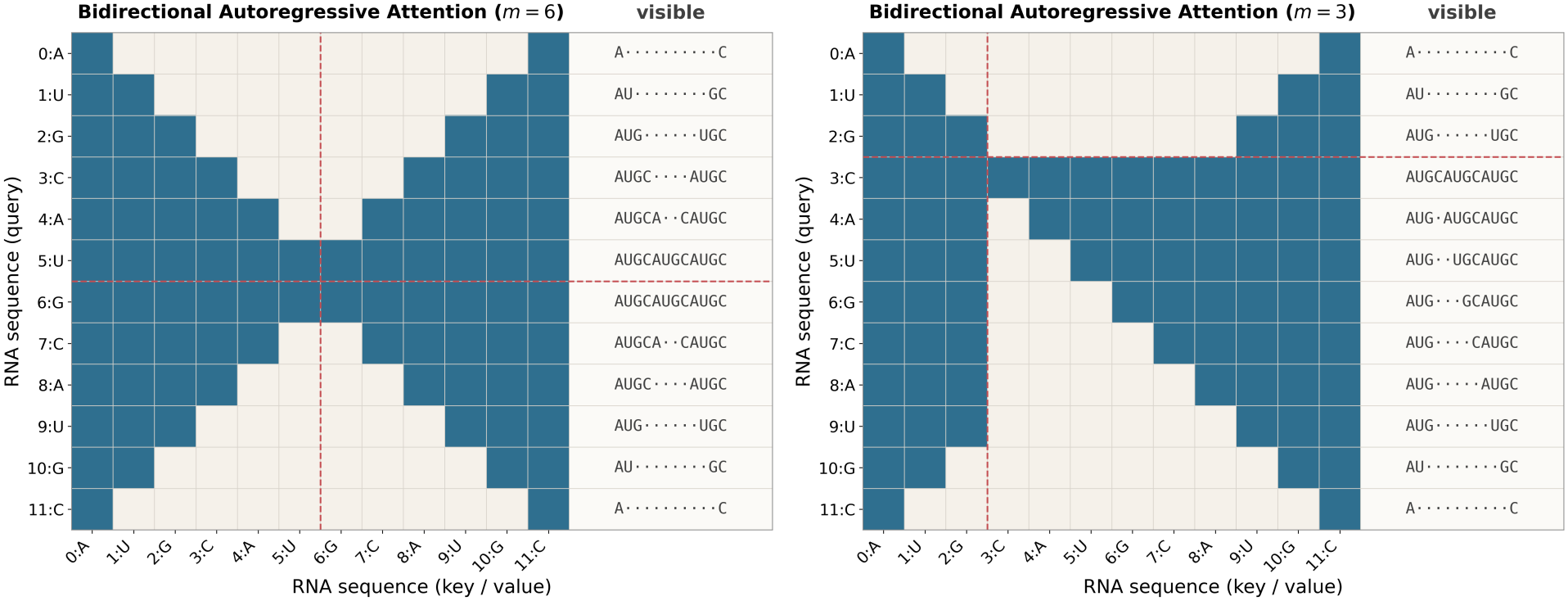
Visualization of bidirectional autoregressive attention for a sequence of length *T* = 12 with merge indices *m* = 6 (left) and *m* = 3 (right). The dashed red line marks the merge index.

### 2.3 Model Architecture

As illustrated in Figure 1, B-RANKO uses a decoder-only Transformer architecture [24] adapted for bidirectional autoregressive RNA modeling. In the configuration used in this work, the model operates on nucleotide-level tokens with a vocabulary of size 8, including the special tokens `<PAD>`, `<SOS>`, `<EOS>`, and `<UNK>`, and the nucleotide tokens `A`, `C`, `G`, and `U` (with `T` mapped to `U`). Tokens are embedded into a 640-dimensional space and processed by a stack of 30 Transformer blocks with 20 attention heads per block. Each block applies masked multi-head self-attention followed by a feed-forward transition layer, and the model uses rotary positional embeddings (RoPE [25]) to encode token order.

Unlike standard causal decoders, B-RANKO replaces the causal mask with the bidirectional autoregressive attention mask described in Section 2.2.

### 2.4 Training Objective

The model is trained to perform autoregressive next-token prediction from both sequence ends toward the merge index *m*. Given an input sequence **x** = (*x*_0_, …, *x*_*T −*1_), the left output head predicts the next token in the left-to-right direction, 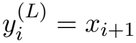, and the right output head predicts the next token in the right-to-left direction,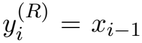. During training, *m* is sampled uniformly from {1, …, *T –* 1} for each training example, so an interior position is predicted by the left head in some iterations and by the right head in others. We expect this to encourage the shared backbone to encode context from both directions.

Positions *i < m* are supervised by the left head and positions *i ≥ m* by the right head, giving the training objective

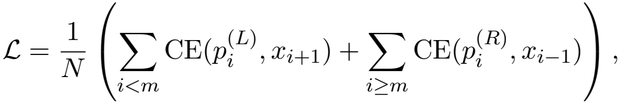

where CE denotes token-level cross-entropy and *N* is the number of valid supervised positions. B-RANKO is pretrained on sequences of 16–1024 nt from RNAcentral [26], Rfam [27] and GARNET [28]. Dataset construction and training hyperparameters are given in Supplementary A.2.

### 2.5 Inference and Representation Extraction

In the generative setting, B-RANKO performs fixed-length inward decoding. A target length is sampled from an empirical length distribution, the sequence is initialized with `<SOS>` and `<EOS>`, and tokens are sampled at temperature 1.0 from both ends toward the center, *m* = *⌊T/*2*⌋*. If the fronts meet at a single position, the left and right logits are averaged before sampling it.

For representation extraction, complete sequences are encoded with full bidirectional attention (Supplementary C.3). Sequence-level embeddings use covariance pooling [29] (Supplementary C.2),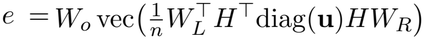, where *H ∈ ℝ*^*T ×d*^ are the final-layer token states, **u** marks the *n* nucleotide positions, and 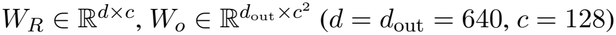

## 3 Results

### 3.1 RNA Sequence Generation

We first evaluate RNA sequence generation by asking whether the generated sequences look like realistic RNAs from the target distribution rather than arbitrary strings. For this analysis, we compare B-RANKO with RANKO and GenerRNA using sequence length, GC content, average minimum free energy (AMFE), novelty and diversity. Sequence length, GC content and AMFE measure whether generated RNAs match the compositional and thermodynamic properties of natural sequences, with GC content and AMFE serving as proxies for folding plausibility. Novelty and diversity measure how the samples relate to the reference set and to each other. For novelty, each generated sequence is classified by its best MMseqs2 hit in the reference set as *known* (strong match), *partially novel* (weaker homology) or *novel* (no sufficiently strong hit). A model that fits the data well should produce all three: known sequences show that it has learned real RNA families, partially novel sequences that it generates variants of them, and novel sequences that it moves beyond the training examples. Diversity is measured by clustering the samples with MMseqs2. The diversity ratio, the number of clusters divided by the number of samples, indicates how broadly they cover sequence space. For both novelty and diversity, we use the B-RANKO pretraining set as the reference. Although this corresponds only to the shorter-sequence subset of the full RNAcentral data used to train RANKO and GenerRNA, which may favor those models on these metrics, it provides the fairest common basis for evaluation. B-RANKO’s sample lengths are drawn from an empirical length distribution (Section 2.5), so its length distribution is set by construction rather than learned. Metric definitions are provided in Supplementary A.3.

All three models produce thermodynamically plausible RNA sequences, with GC content and AMFE distributions similar to the reference set (Figure 3). The clearest differences appear in novelty and diversity. Compared with GenerRNA, both B-RANKO and RANKO generate more novel sequences distributed across a larger number of clusters (Table 1). Between the two, RANKO shows slightly higher novelty. Both have diversity ratios above that of the reference set (0.815 and 0.789 vs. 0.694).

**Table 1:** MMseqs2 clustering diversity of samples from the pretrained models, 10,000 sequences per model and for the reference set.

| Model | Clusters | Diversity Ratio | Exact Duplicate Rate |
|---|---|---|---|
| B-RANKO | 8149 | 0.815 | 0.0% |
| RANKO | 7894 | 0.789 | 0.1% |
| GenerRNA | 4009 | 0.401 | 0.1% |
| Reference | 6942 | 0.694 | 0.0% |

**Figure 3:**
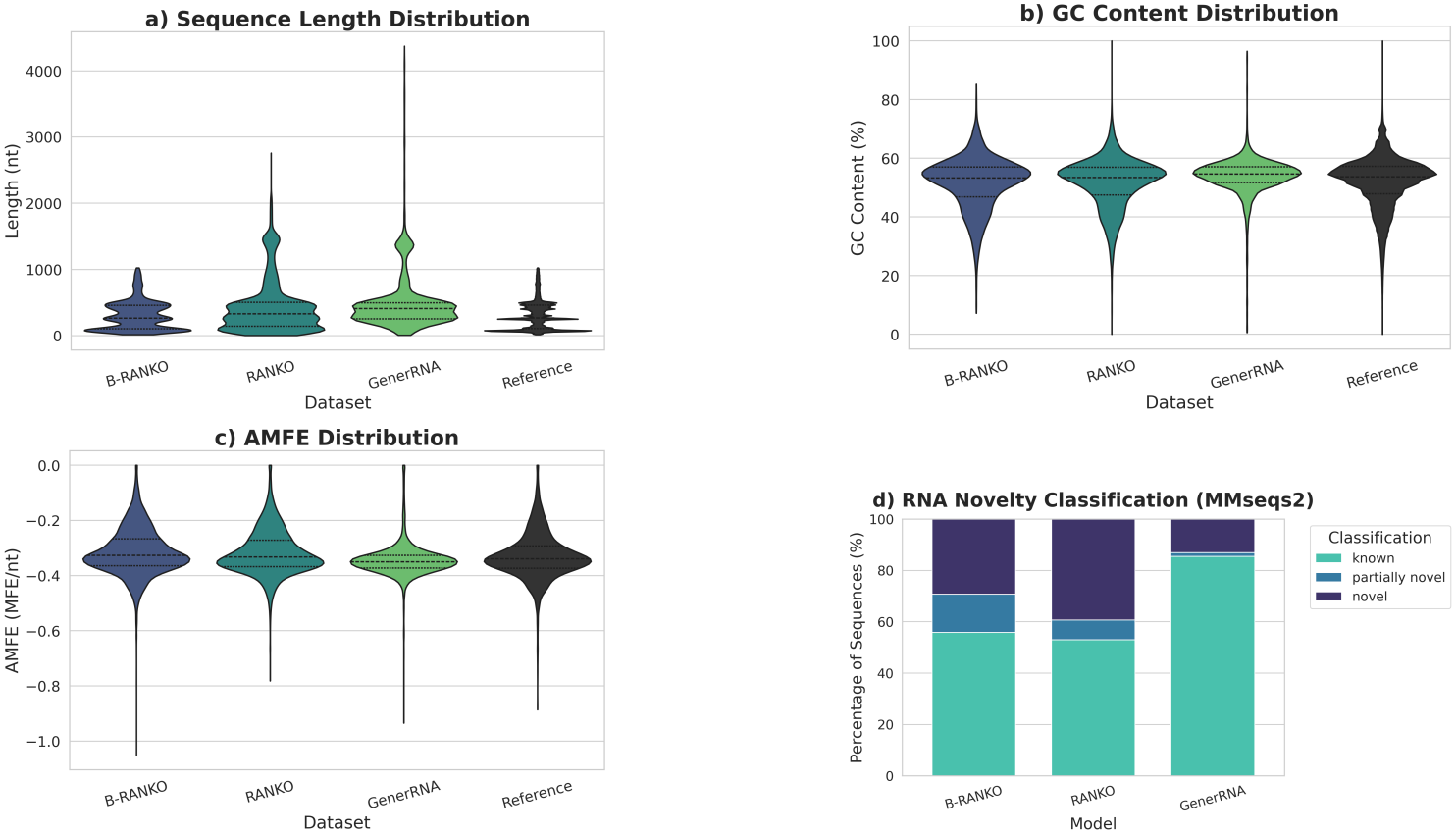
Samples from the pretrained models compared with the reference set.

**Figure 4:**
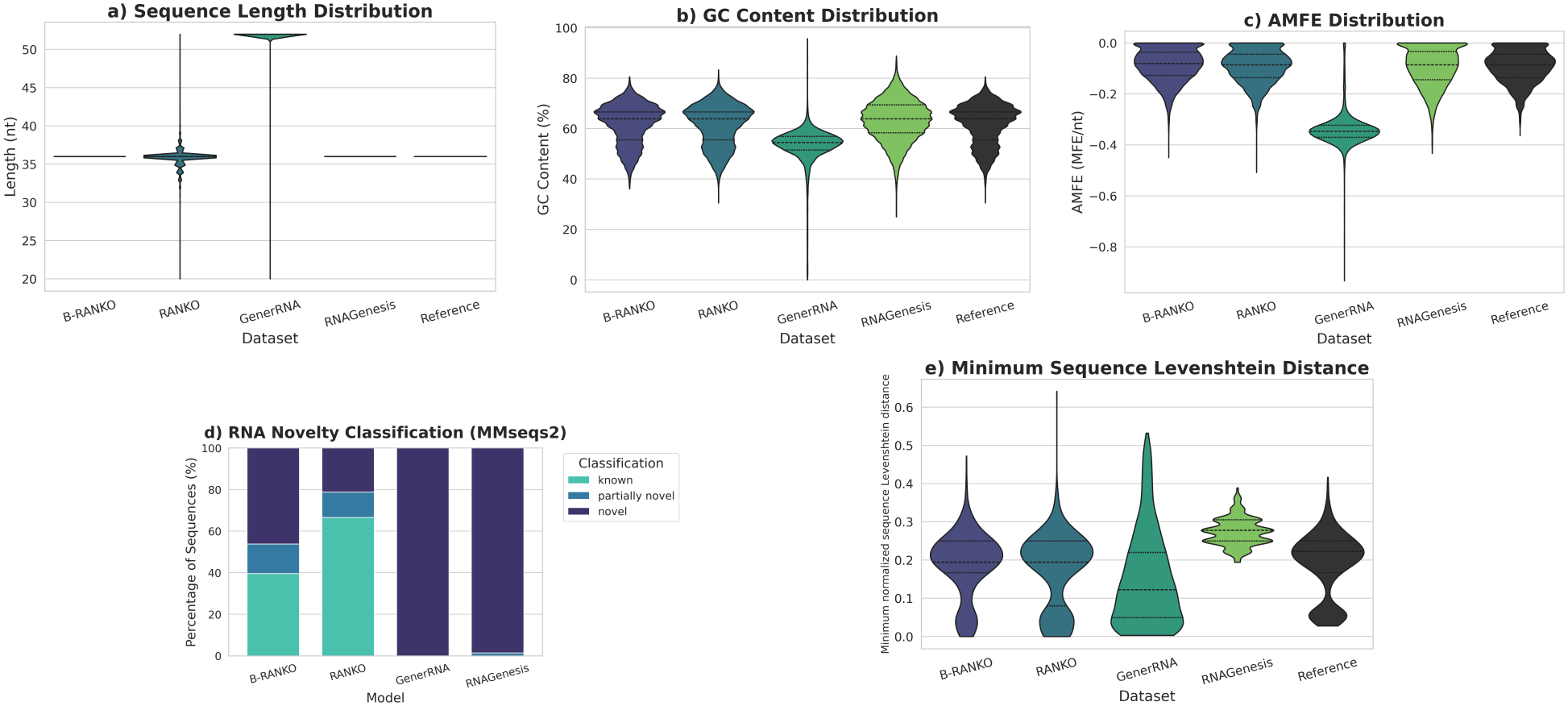
Samples after fine-tuning on AptaDiff Dataset A.

### 3.2 Aptamer Design

While the previous section examined whether the models reproduce the global statistics of natural RNA sequences, we now evaluate their transfer to a downstream design task. Specifically, we assess aptamer generation on AptaDiff [30] Dataset A, an HT-SELEX benchmark of fixed-length 36-nt RNA sequences targeting IGFBP3, using the same metrics as in the pretraining analysis reported in Section 3.1 together with minimum sequence Levenshtein distance. For each generated sequence, this is the normalized edit distance to its closest neighbor among the other generated sequences, so low values indicate near-duplicates. We use it only here because all target sequences have the same length, so the distance reflects sequence variation rather than differences in length. We additionally include RNAGenesis only in this downstream comparison, as the authors provide a publicly available fine-tuned checkpoint but not a comparable pretrained checkpoint. They report [17] pretraining the Q-former and decoder on top of a frozen encoder for 2 epochs on the ncRNA subset of the Ensembl database (1.1 million sequences), followed by fine-tuning on AptaDiff Dataset A for 10 epochs. B-RANKO and RANKO were fine-tuned on AptaDiff Dataset A for 10 epochs with a learning rate of 10^*−*5^, while GenerRNA was fine-tuned for 30 epochs using the authors’ instructions on Hugging Face.

Under this setup, GenerRNA shows the weakest adaptation to Dataset A. Its samples vary in length and differ from the target aptamers in GC content and AMFE, appearing closer to its pretraining distribution. B-RANKO’s and RNAGenesis’s lengths are fixed at 36 nt by construction.

B-RANKO, RANKO and RNAGenesis match the GC content and AMFE of the reference more closely. Of these, B-RANKO is closest to the reference in diversity (Table 2), while RANKO is more conservative, with more *known* sequences and more exact duplicates. RNAGenesis samples are almost all novel and almost all distinct, more spread out than the target set itself. This suggests that they remain partly shaped by its pretraining data, although we cannot check this directly because that data is not publicly available.

**Table 2:** MMseqs2 clustering diversity on AptaDiff Dataset A, 10,000 sequences per model and for the reference set.

| Model | Clusters | Diversity Ratio | Exact Duplicate Rate |
|---|---|---|---|
| B-RANKO | 7335 | 0.734 | 12.5% |
| RANKO | 5609 | 0.561 | 27.5% |
| GenerRNA | 4067 | 0.407 | 0.1% |
| RNAGenesis | 9991 | 0.999 | 0.0% |
| Reference | 7169 | 0.717 | 0.0% |

### 3.3 Representation Learning on BEACON

To assess the quality and transferability of the learned representations, we evaluate them on the 13 BEACON tasks, grouped into three categories. The *structure* tasks are secondary structure (SSP), contact map (CMP) and distance map (DMP) prediction, and structural score imputation (SSI). The *function* tasks are splice site (SPL), APA isoform (APA), non-coding RNA function (ncRNA) and modification (Modif) prediction, and mean ribosome loading (MRL). The *engineering* tasks are vaccine degradation prediction (VDP), programmable RNA switches (PRS), and CRISPR on- and off-target prediction (CRI-On, CRI-Off). Further details on the datasets and evaluation protocols are provided in the BEACON paper [19], while the task heads and hyperparameter settings used in our experiments are summarized in Supplementary A.4.

We compare B-RANKO against the full set of pretrained RNA language model baselines reported in the BEACON benchmark paper [19], and additionally include strong RNA foundation models with publicly available BEACON results. Specifically, RNAGenesis results are taken from [17], while RiNALMo and EvoFlow-RNA are taken from [16]. RANKO is evaluated only on the sequence-level tasks. The nucleotide-level tasks need a representation of each nucleotide in whole-sequence context, which RANKO cannot provide. BPE tokens do not map one-to-one to nucleotides, and its causal attention limits each position to preceding context.

Overall, B-RANKO is competitive but uneven across BEACON. It performs strongest on the engineering benchmarks, achieving the best results on vaccine degradation and CRISPR off-target prediction and ranking in the top three on programmable RNA switches. It also reaches second-best performance on distance map prediction and top-three performance on Non-coding RNA Function Classification (ncRNA), but lags the strongest baselines on some structure tasks, especially secondary structure prediction and contact map prediction. B-RANKO transfers particularly well to engineering and some sequence-level prediction tasks, while its gains are less consistent for finer-grained local structural predictions. At the same time, its strong distance map performance indicates that it still captures structurally relevant information, even if this is not reflected uniformly across all structure benchmarks.

### 3.4 Ablation: Bidirectional versus causal attention at matched scale

In Table 3, B-RANKO outperforms RANKO on all seven tasks on which both are evaluated, despite being about four times smaller. The two models, however, also differ in tokenization and size, so this comparison does not isolate the effect of the attention mask. To isolate it, we pretrain two 33M-parameter models that differ only in the mask, one with bidirectional autoregressive attention and one with causal attention. Both use single-nucleotide tokens and are trained for 10 epochs on Rfam representatives with identical optimization settings. We evaluate both on all 13 BEACON tasks with mean pooling, extracting representations with full attention and with the mask each model was pretrained with (complete configuration and hyperparameters are in Supplementary C.4).

**Table 3:** Performance on the 13 BEACON downstream tasks. Best, second-best, and third-best results in each metric are highlighted. For B-RANKO and RANKO, mean_(*sd*)_ over three runs are reported.

| Model | Size | SSP<br>F1 (%) | CMP<br>P@L (%) | DMP<br>R<sup>2</sup> (%) | SSI<br>R<sup>2</sup> (%) | SPL<br>ACC@K (%) | APA<br>R<sup>2</sup> (%) | NcRNA<br>ACC (%) | Modif<br>AUC (%) | MRL<br>R<sup>2</sup> (%) | VDP<br>MCRMSE↓ | PRS<br>R<sup>2</sup> (%) | CRI-On<br>SC (%) | CRI-Off<br>SC (%) |
|---|---|---|---|---|---|---|---|---|---|---|---|---|---|---|
| RNA-FM | 99.52M | 68.50 | 47.56 | 51.45 | 42.36 | 34.84 | 70.32 | 96.81 | 94.98 | 79.47 | 0.347 | 55.98 | 31.62 | 2.49 |
| RNABERT | 0.48M | 57.27 | 45.21 | 48.19 | 31.62 | 0.18 | 57.66 | 68.95 | 82.82 | 29.79 | 0.378 | 54.60 | 29.77 | 4.27 |
| RNA-MSM | 95.92M | 57.98 | 57.26 | 37.49 | 39.22 | 38.33 | 70.40 | 84.85 | 94.89 | 83.48 | 0.330 | 56.94 | 34.92 | 3.85 |
| Splice-H510 | 19.45M | 64.93 | 45.80 | 55.56 | 38.91 | 44.80 | 58.65 | 95.92 | 62.57 | 83.49 | 0.321 | 54.90 | 26.61 | 4.00 |
| Splice-MS510 | 19.45M | 43.24 | 52.64 | 10.27 | 38.58 | 50.55 | 52.46 | 95.87 | 55.87 | 84.98 | 0.315 | 50.98 | 27.13 | 3.49 |
| Splice-MS1024 | 19.72M | 68.26 | 47.32 | 55.89 | 39.22 | 48.52 | 60.03 | 96.05 | 53.45 | 67.15 | 0.313 | 57.72 | 27.59 | 5.00 |
| UTR-LM-MRL | 1.21M | 59.71 | 45.51 | 55.21 | 39.52 | 36.20 | 64.99 | 89.97 | 56.41 | 77.78 | 0.325 | 57.28 | 28.49 | 4.28 |
| UTR-LM-TE&EL | 1.21M | 59.57 | 60.32 | 54.94 | 40.15 | 37.35 | 72.09 | 81.33 | 59.70 | 82.50 | 0.319 | 53.37 | 32.49 | 2.91 |
| UTRBERT-3mer | 86.14M | 60.37 | 51.03 | 50.95 | 34.31 | 44.24 | 69.52 | 92.88 | 95.14 | 83.89 | 0.337 | 56.83 | 29.92 | 4.48 |
| UTRBERT-4mer | 86.53M | 59.41 | 44.91 | 47.77 | 33.22 | 42.04 | 72.71 | 94.32 | 95.10 | 82.90 | 0.341 | 56.43 | 23.20 | 3.11 |
| UTRBERT-5mer | 86.45M | 47.92 | 44.71 | 48.67 | 31.27 | 39.19 | 72.70 | 93.04 | 94.78 | 75.64 | 0.343 | 57.16 | 25.74 | 3.93 |
| UTRBERT-6mer | 98.05M | 38.56 | 51.56 | 50.02 | 29.93 | 38.58 | 71.17 | 93.12 | 95.08 | 83.60 | 0.340 | 57.14 | 28.60 | 4.90 |
| BEACON-B | 87M | 64.18 | 60.81 | 56.28 | 38.78 | 37.43 | 70.59 | 94.63 | 94.74 | 72.29 | 0.320 | 54.67 | 26.01 | 4.42 |
| BEACON-B512 | 87M | 58.75 | 61.20 | 56.82 | 39.13 | 37.24 | 72.00 | 94.99 | 94.92 | 72.35 | 0.320 | 55.20 | 28.17 | 3.82 |
| RNAGenesis | 866M | 73.72 | 71.16 | 70.31 | – | 37.70 | 89.15 | 97.81 | 95.14 | 85.75 | – | 58.12 | 35.55 | – |
| RiNALMo (mega) | 150M | 75.02 | 74.03 | 53.11 | 42.70 | 38.49 | 81.03 | 91.89 | 95.04 | 85.91 | 0.281 | 56.02 | 40.99 | 5.44 |
| EvoFlow-RNA | 150M | 74.51 | 74.05 | 47.72 | 42.52 | 32.66 | 72.61 | 86.46 | 71.06 | 86.28 | 0.47 | 55.03 | 41.48 | 5.34 |
| RANKO (ablation) | 651.83M | – | – | – | – | – | 60.49<sub>(1.07)</sub> | 91.44<sub>(0.12)</sub> | 85.69<sub>(0.05)</sub> | 68.53<sub>(0.41)</sub> | – | 48.39<sub>(0.37)</sub> | 18.79<sub>(1.23)</sub> | 7.46<sub>(1.40)</sub> |
| B-RANKO | 150M | 49.12<sub>(0.26)</sub> | 44.66<sub>(0.98)</sub> | 57.78<sub>(0.52)</sub> | 39.69<sub>(0.36)</sub> | 39.38<sub>(1.10)</sub> | 69.11<sub>(1.00)</sub> | 96.72<sub>(0.59)</sub> | 95.08<sub>(0.11)</sub> | 85.04<sub>(0.25)</sub> | 0.264<sub>(0.003)</sub> | 57.67<sub>(0.16)</sub> | 26.27<sub>(6.48)</sub> | 10.77<sub>(1.59)</sub> |

**Table 4:** Matched-scale comparison on BEACON. Mean_(sd)_ over three seeds.

|  | Bidirectional |  | Causal |  |
|---|---|---|---|---|
| Task | Full | Inward | Causal | Full |
| SSP [F1 %] | 41.68<sub>(1.49)</sub> | 42.98<sub>(1.08)</sub> | 39.41<sub>(2.01)</sub> | 42.06<sub>(1.54)</sub> |
| CMP [P@L %] | 41.83<sub>(1.47)</sub> | 43.02<sub>(0.45)</sub> | 33.79<sub>(0.85)</sub> | 38.54<sub>(0.64)</sub> |
| DMP [R<sup>2</sup> %] | 56.63<sub>(0.60)</sub> | 55.94<sub>(0.70)</sub> | 52.13<sub>(0.57)</sub> | 54.76<sub>(0.13)</sub> |
| SSI [R<sup>2</sup> %] | 39.25<sub>(0.09)</sub> | 20.96<sub>(0.15)</sub> | 8.90<sub>(0.11)</sub> | 38.88<sub>(0.25)</sub> |
| SpliceAI [ACC@K %] | 37.97<sub>(1.07)</sub> | 18.86<sub>(0.97)</sub> | 12.28<sub>(1.59)</sub> | 37.91<sub>(0.55)</sub> |
| APA [R<sup>2</sup> %] | 69.67<sub>(0.63)</sub> | 61.55<sub>(2.06)</sub> | 64.69<sub>(3.42)</sub> | 69.37<sub>(0.73)</sub> |
| NcRNA [ACC %] | 94.27<sub>(0.85)</sub> | 92.69<sub>(0.60)</sub> | 91.74<sub>(0.37)</sub> | 93.37<sub>(0.47)</sub> |
| Modif [AUC %] | 94.89<sub>(0.12)</sub> | 73.22<sub>(0.10)</sub> | 94.56<sub>(0.11)</sub> | 94.76<sub>(0.05)</sub> |
| MRL [R<sup>2</sup> %] | 82.57<sub>(0.23)</sub> | 72.22<sub>(0.85)</sub> | 72.80<sub>(0.48)</sub> | 79.70<sub>(0.67)</sub> |
| VDP [MCRMSE ↓] | 0.269<sub>(0.002)</sub> | 0.349<sub>(0.002)</sub> | 0.360<sub>(0.002)</sub> | 0.283<sub>(0.001)</sub> |
| PRS [R<sup>2</sup> %] | 57.10<sub>(0.05)</sub> | 57.02<sub>(0.18)</sub> | 55.89<sub>(0.19)</sub> | 56.00<sub>(0.22)</sub> |
| CRI-On [SC %] | 28.27<sub>(4.73)</sub> | 33.30<sub>(3.36)</sub> | 27.49<sub>(2.64)</sub> | 27.55<sub>(3.41)</sub> |
| CRI-Off [SC %] | 11.18<sub>(0.92)</sub> | 11.07<sub>(0.76)</sub> | 8.55<sub>(1.65)</sub> | 10.60<sub>(0.88)</sub> |

With each model used as intended, the bidirectional model with full-attention extraction outperforms the causal model on all 13 tasks, by +1.21 on programmable RNA switches up to +30.35 on structural score imputation. Most of this gap comes from extraction: with full attention, the causal model recovers much of it, raising its structural score imputation from 8.90 to 38.88. With extraction fixed to full attention, the bidirectional model is still better on 12 of 13 tasks, five of them by more than two pooled standard deviations. Supplementary C reports two further ablations on the 150M B-RANKO: full-attention versus inward extraction (Supplementary C.3) and covariance versus mean pooling (Supplementary C.2).

## 4 Discussion

B-RANKO shows that a decoder-only ncRNA foundation model can support both sequence generation and transferable representation learning within a single architecture. In pretrained inference, it generates RNA sequences with realistic GC-content and AMFE distributions while producing a large share of novel sequences that fall into more clusters than the reference set. After fine-tuning, it adapts effectively to aptamer generation, producing sequences that remain well aligned with the target distribution without becoming overly conservative or repetitive. On BEACON, its strongest results are on RNA engineering tasks: among the compared pretrained models it has the lowest vaccine degradation error and the highest CRISPR off-target score, and it ranks third on programmable RNA switches (PRS). At matched scale, bidirectional autoregressive pretraining gives better representations than causal pretraining, although most of the difference comes from extracting representations with full attention (Section 3.4). Together, these findings support the proposed bidirectional autoregressive attention mechanism and show that B-RANKO can unify generation and downstream transfer in a single decoder-only model.

At the same time, the results also point to clear directions for future work. Performance is not uniformly strong across all representation-learning tasks, with weaker results on secondary structure prediction and contact map prediction than on distance map prediction and engineering benchmarks. This suggests that B-RANKO captures useful global structural and sequence-level signals, but may not yet learn sufficiently fine-grained nucleotide-level embeddings for all structure tasks. Scaling the model and pretraining setup is therefore a natural next step, particularly for tasks that depend on precise local structural representations.

B-RANKO’s two roles also suggest a direct next step: using its representations to evaluate its own samples. Prediction heads trained on B-RANKO embeddings, such as those for vaccine degradation or CRISPR off-target activity, can score generated sequences, and these scores can steer generation, either by filtering samples or as rewards in reinforcement learning with verifiable rewards.

Finally, the generated sequences have so far been evaluated only in silico, and their function remains to be tested experimentally. Aptamer design is a natural first test, since B-RANKO already adapts to a target family: generated candidates could be synthesized and tested for binding, or used to seed SELEX libraries.

## 5 Conclusion

In this work, we introduced **B-RANKO**, a 150M-parameter bidirectional decoder-only ncRNA foundation model that unifies *de novo* sequence generation and transferable representation learning within a single architecture. More broadly, B-RANKO shows that extending the standard causal autoregressive attention used in RANKO with bidirectional autoregressive attention can unify generation and downstream transfer within a single decoder-only RNA model. To enable this, we proposed a novel bidirectional autoregressive attention mechanism that supports decoder-only training over both forward and reverse sequence directions while allowing both native sequence generation and embedding extraction for downstream tasks. In pretrained inference, B-RANKO generates realistic RNA sequences and, as assessed by MMseqs2, produces novel sequences that extend beyond the pretraining corpus. After fine-tuning, it adapts efficiently to task-specific generation, illustrated here on aptamer design. On BEACON, it learns useful transferable representations and is particularly strong on RNA engineering tasks, with the lowest vaccine degradation error among the compared pretrained models. Overall, these findings establish bidirectional decoder-only modeling as a viable direction for RNA foundation models and show that a single model can support generation, efficient adaptation, and downstream prediction within one unified framework.

## Acknowledgments

This work has been supported by the Croatian Science Foundation under the project “Deep Learning-Based RNA Tertiary Structure Prediction and Generation” (IP-2024-05-1554). This research was performed using the advanced computing service provided by the University of Zagreb University Computing Centre (SRCE).

## References

[1] Francis H. C. Crick. Central dogma of molecular biology. Nature, 227(5258):561–563, 1970. doi: 10.1038/227561a0. URL https://doi.org/10.1038/227561a0.

[2] Jennifer A. Doudna and Thomas R. Cech. The chemical repertoire of natural ribozymes. Nature, 418 (6894):222–228, 2002. doi: 10.1038/418222a. URL https://doi.org/10.1038/418222a.

[3] David P. Bartel. Micrornas: genomics, biogenesis, mechanism, and function. Cell, 116(2):281–297, 2004. doi: 10.1016/S0092-8674(04)00045-5. URL https://doi.org/10.1016/S0092-8674(04)00045-5.

[4] Andrew Fire, SiQun Xu, Mary K. Montgomery, Samuel A. Kostas, Stephen E. Driver, and Craig C. Mello. Potent and specific genetic interference by double-stranded rna in C. elegans. Nature, 391 (6669):806–811, 1998. doi: 10.1038/35888. URL https://doi.org/10.1038/35888.

[5] John L. Rinn and Howard Y. Chang. Genome regulation by long noncoding RNAs. Annual Review of Biochemistry, 81:145–166, 2012. doi: 10.1146/annurev-biochem-051410-092902. URL https://doi.org/10.1146/annurev-biochem-051410-092902.

[6] Martin Jinek, Krzysztof Chylinski, Ines Fonfara, Michael Hauer, Jennifer A. Doudna, and Emmanuelle Charpentier. A programmable dual-RNA-guided DNA endonuclease in adaptive bacterial immunity. Science, 337(6096):816–821, 2012. doi: 10.1126/science.1225829. URL https://doi.org/10.1126/science.1225829.

[7] Andrew D. Ellington and Jack W. Szostak. In vitro selection of RNA molecules that bind specific ligands. Nature, 346(6287):818–822, 1990. doi: 10.1038/346818a0. URL https://doi.org/10.1038/346818a0.

[8] Ugur Sahin, Katalin Karikó, and Özlem Türeci. mRNA-based therapeutics—developing a new class of drugs. Nature Reviews Drug Discovery, 13(10):759–780, 2014. doi: 10.1038/nrd4278. URL https://doi.org/10.1038/nrd4278.

[9] Norbert Pardi, Michael J. Hogan, Frederick W. Porter, and Drew Weissman. mRNA vaccines — a new era in vaccinology. Nature Reviews Drug Discovery, 17(4):261–279, 2018. doi: 10.1038/nrd.2017.243. URL https://doi.org/10.1038/nrd.2017.243.

[10] John Jumper, Richard Evans, Alexander Pritzel, Tim Green, Michael Figurnov, Olaf Ronneberger, Kathryn Tunyasuvunakool, Russ Bates, Augustin Zídek, Anna Potapenko, Alex Bridgland, Clemens Meyer, Simon A. A. Kohl, Andrew J. Ballard, Andrew Cowie, Bernardino Romera-Paredes, Stanislav Nikolov, Rishub Jain, Jonas Adler, Trevor Back, Stig Petersen, David Reiman, Ellen Clancy, Michal Zielinski, Martin Steinegger, Michalina Pacholska, Tamas Berghammer, Sebastian Bodenstein, David Silver, Oriol Vinyals, Andrew W. Senior, Koray Kavukcuoglu, Pushmeet Kohli, and Demis Hassabis. Highly accurate protein structure prediction with alphafold. Nature, 596:583–589, 2021. doi: 10.1038/s41586-021-03819-2. URL https://www.nature.com/articles/s41586-021-03819-2.

[11] Garyk Brixi, Matthew G. Durrant, Jerome Ku, Mohsen Naghipourfar, Michael Poli, Gwanggyu Sun, Greg Brockman, Daniel Chang, Alison Fanton, Gabriel A. Gonzalez, Samuel H. King, David B. Li, Aditi T. Merchant, Eric Nguyen, Chiara Ricci-Tam, David W. Romero, Jonathan C. Schmok, Ali Taghibakhshi, Anton Vorontsov, Brandon Yang, et al. Genome modelling and design across all domains of life with evo 2. Nature, 652:1349–1361, 2026. doi: 10.1038/s41586-026-10176-5. URL https://www.nature.com/articles/s41586-026-10176-5.

[12] Jiayang Chen, Zhihang Hu, Siqi Sun, Qingxiong Tan, Yixuan Wang, Qinze Yu, Licheng Zong, Liang Hong, Jin Xiao, Tao Shen, Irwin King, and Yu Li. Interpretable rna foundation model from unannotated data for highly accurate rna structure and function predictions. arXiv preprint arXiv:2204.00300, 2022. URL https://arxiv.org/abs/2204.00300.

[13] Rafael Josip Penić, Tin Vlašić, Roland G. Huber, Yue Wan, and Mile Šikić. Rinalmo: general-purpose rna language models can generalize well on structure prediction tasks. Nature Communications, 16:5671, 2025. doi: 10.1038/s41467-025-60872-5. URL https://www.nature.com/articles/s41467-025-60872-5.

[14] Shuxian Zou, Tianhua Tao, Sazan Mahbub, Caleb N. Ellington, Robin Algayres, Dian Li, Yonghao Zhuang, Hongyi Wang, L. Song, and Eric P. Xing. A large-scale foundation model for rna function and structure prediction. bioRxiv, 2024. doi: 10.1101/2024.11.28.625345. URL https://doi.org/10.1101/2024.11.28.625345.

[15] Yichong Zhao, Kenta Oono, Hiroki Takizawa, and Masaaki Kotera. GenerRNA: A generative pre-trained language model for de novo RNA design. bioRxiv, 2024. doi: 10.1101/2024.02.01.578496. URL http://biorxiv.org/lookup/doi/10.1101/2024.02.01.578496.

[16] Sawan Patel, Fred Zhangzhi Peng, Keith Fraser, Adam D. Friedman, Pranam Chatterjee, and Sherwood Yao. EvoFlow-RNA: Generating and representing non-coding RNA with a language model. bioRxiv, 2025. doi: 10.1101/2025.02.25.639942. URL http://biorxiv.org/lookup/doi/10.1101/2025.02.25.639942.

[17] Zaixi Zhang, Linlin Chao, Ruofan Jin, Yikun Zhang, Guowei Zhou, Yujie Yang, Yukang Yang, Kaixuan Huang, Qirong Yang, Ziyao Xu, Xiaoming Zhang, L. Cong, and Mengdi Wang. Rnagenesis: Foundation model for enhanced rna sequence generation and structural insights. bioRxiv, 2024. doi: 10.1101/2024.12.30.630826. URL https://doi.org/10.1101/2024.12.30.630826.

[18] Martin Steinegger and Johannes Söding. MMseqs2 enables sensitive protein sequence searching for the analysis of massive data sets. Nature Biotechnology, 35(11):1026–1028, 2017. doi: 10.1038/nbt.3988. URL https://doi.org/10.1038/nbt.3988.

[19] Yuchen Ren, Zhiyuan Chen, Lifeng Qiao, Hongtai Jing, Yuchen Cai, Sheng Xu, Peng Ye, Xinzhu Ma, Siqi Sun, Hongliang Yan, Dong Yuan, Wanli Ouyang, and Xihui Liu. Beacon: Benchmark for comprehensive rna tasks and language models. arXiv preprint arXiv:2406.10391, 2024. URL https://arxiv.org/abs/2406.10391.

[20] Aron M. Yoffe, Peter Prinsen, William M. Gelbart, and Avinoam Ben-Shaul. The ends of a large RNA molecule are necessarily close. Nucleic Acids Research, 39(1):292–299, 2011.

[21] Zhilin Yang, Zihang Dai, Yiming Yang, Jaime Carbonell, Ruslan Salakhutdinov, and Quoc V. Le. Xlnet: Generalized autoregressive pretraining for language understanding. In Advances in Neural Information Processing Systems, volume 32, 2019.

[22] Anh Nguyen, Nikos Karampatziakis, and Weizhu Chen. Meet in the middle: A new pre-training paradigm. In Advances in Neural Information Processing Systems, volume 36, 2023.

[23] Roman Klypa, Alberto Bietti, and Sergei Grudinin. Bang: Bidirectional anchored generation for conditional rna design. In Proceedings of the 42nd International Conference on Machine Learning, volume 267 of Proceedings of Machine Learning Research, 2025.

[24] Ashish Vaswani, Noam Shazeer, Niki Parmar, Jakob Uszkoreit, Llion Jones, Aidan N. Gomez, Łukasz Kaiser, and Illia Polosukhin. Attention is all you need. In Advances in Neural Information Processing Systems, 2017.

[25] Jianlin Su, Yu Lu, Shengfeng Pan, Bo Wen, and Yunfeng Liu. Roformer: Enhanced transformer with rotary position embedding. arXiv preprint arXiv:2104.09864, 2021. URL https://arxiv.org/abs/2104.09864.

[26] The RNAcentral Consortium. RNAcentral 2021: secondary structure integration, improved sequence search and new member databases. Nucleic Acids Research, 49(D1):D212–D220, 2021. doi: 10.1093/nar/gkaa921. URL https://doi.org/10.1093/nar/gkaa921.

[27] Ioanna Kalvari, Eric P. Nawrocki, Nancy Ontiveros-Palacios, Joanna Argasinska, Kevin Lamkiewicz, Manja Marz, Sam Griffiths-Jones, Claire Toffano-Nioche, Daniel Gautheret, Zasha Weinberg, Elena Rivas, Sean R. Eddy, Robert D. Finn, Alex Bateman, and Anton I. Petrov. Rfam 14: expanded coverage of metagenomic, viral and microRNA families. Nucleic Acids Research, 49(D1):D192–D200, 2021. doi: 10.1093/nar/gkaa1047. URL https://doi.org/10.1093/nar/gkaa1047.

[28] Marena I. Trinidad and Jamie H. D. Cate. GARNET database, 2024. URL 10.5281/zenodo.14642546. Data retrieved from Zenodo repository.

[29] Thomas Dooms, Nicholas K Wang, and Michael T Pearce. Covariance-based sequence pooling. Goodfire Research, 2026.

[30] Z. Wang et al. Aptadiff: de novo design and optimization of aptamers based on diffusion models. Briefings in Bioinformatics, 25(6):bbae517, 2024. doi: 10.1093/bib/bbae517. URL https://doi.org/10.1093/bib/bbae517.

