## Supplementary file for "B-RANKO: An ncRNA Foundation Model with Bidirectional Decoder-Only Architecture"

**Filip Vučenik**[1] **Rafael Josip Penić**[1] **Mile Šikić**[1,2]
[1]Faculty of Electrical Engineering and Computing, University of Zagreb, Zagreb, Croatia
[2]Genome Institute of Singapore, A*STAR, Singapore, Republic of Singapore
{filip.vucenik, rafael-josip.penic, mile.sikic}@fer.unizg.hr

#### A Hyperparameters and Implementation Details

##### A.1 RANKO: Architecture, Dataset Construction, Preprocessing, and Training Hyperparameters

The pretraining corpus used for RANKO was assembled from the GARNET database [1], Rfam release 15.1 [2], and RNAcentral release 26.0 [3]. All source FASTA files were merged into a single corpus and processed through a uniform cleaning pipeline. Sequences were converted to uppercase, exact duplicates were removed, sequences containing more than 20% ambiguous or non-canonical characters were excluded, and only sequences with lengths between 16 and 2048 nucleotides were retained. This preprocessing yielded a cleaned corpus of 40,054,145 sequences. To obtain a more compact and diverse pretraining set, the cleaned corpus was further clustered with MMseqs2 `easy-linclust` [4] using minimum sequence identity 0.9 and coverage 0.8, producing a representative corpus of 20,020,987 sequences, of which 19,920,987 were used for training and 100,000 for validation.

RANKO used Byte Pair Encoding (BPE) with a vocabulary of 1,024 tokens trained on Rfam. Tokenized sequences were used to train a 650M-parameter decoder-only Transformer with 33 transformer blocks, an embedding dimension of 1,280, and 20 attention heads. FlashAttention was used to improve memory efficiency during long-context training. Training was performed in two stages. In the first stage, the model was trained on the clustered representative corpus. In the second stage, training was resumed from the first-stage checkpoint on the full cleaned dataset, while retaining the representative validation split. Training used 8 NVIDIA H200 GPUs with bf16-mixed precision, per-device batch size 100, and no gradient accumulation, corresponding to a global batch size of 800 sequences per optimizer step. Optimization used AdamW with learning rate $5.0 \times 10^{-5}$, weight decay $0.01$, and $\beta = (0.9, 0.999)$. The learning rate schedule consisted of a 2000-step linear warmup followed by cosine decay to a minimum learning rate of $1.0 \times 10^{-5}$. The cosine schedule was configured for 200,000 iterations in the clustered stage and 150,000 iterations in the full-dataset continuation stage. Gradient clipping with maximum norm 1.0 was applied throughout training, and the model was optimized with standard next-token cross-entropy loss.

##### A.2 B-RANKO: Dataset Construction, Preprocessing, and Training Hyperparameters

B-RANKO uses the same source corpora, cleaning pipeline, and MMseqs2 clustering as RANKO (Supplementary A.1), yielding the same cleaned corpus of 40,054,145 sequences and the same clustered representative set. It differs in the data restriction and training schedule. B-RANKO restricts both training stages to sequences of at most 1024 nucleotides. In the first stage, the model is trained for 4 epochs on the clustered representative dataset, with a validation split of 100,000 sequences held out using a fixed seed of 42. In the second stage, training is resumed from that checkpoint on the full cleaned dataset, which contains 33,640,592 training and 84,046 validation sequences. Each stage required approximately 7 days.

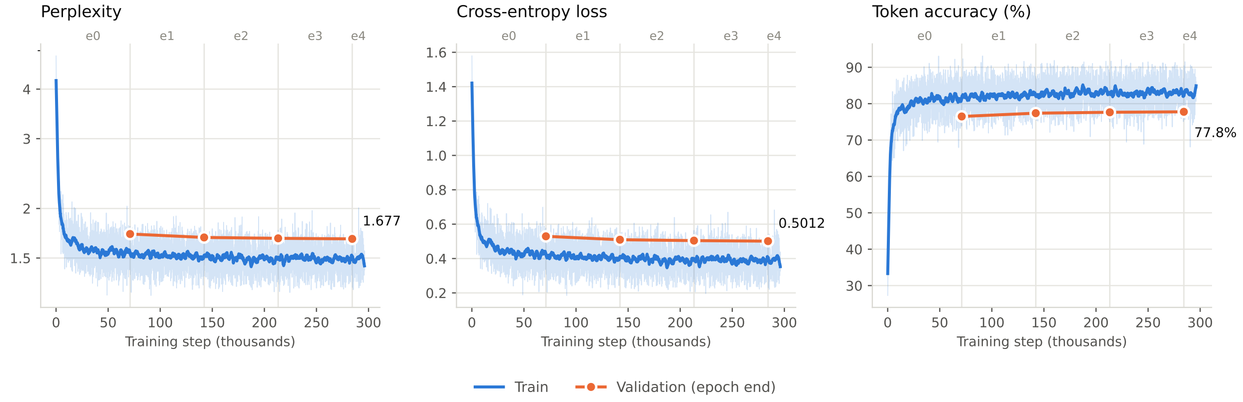


Figure 1: B-RANKO pre-training on the clustered dataset. Markers are epoch-end validation. Perplexity uses a log axis.

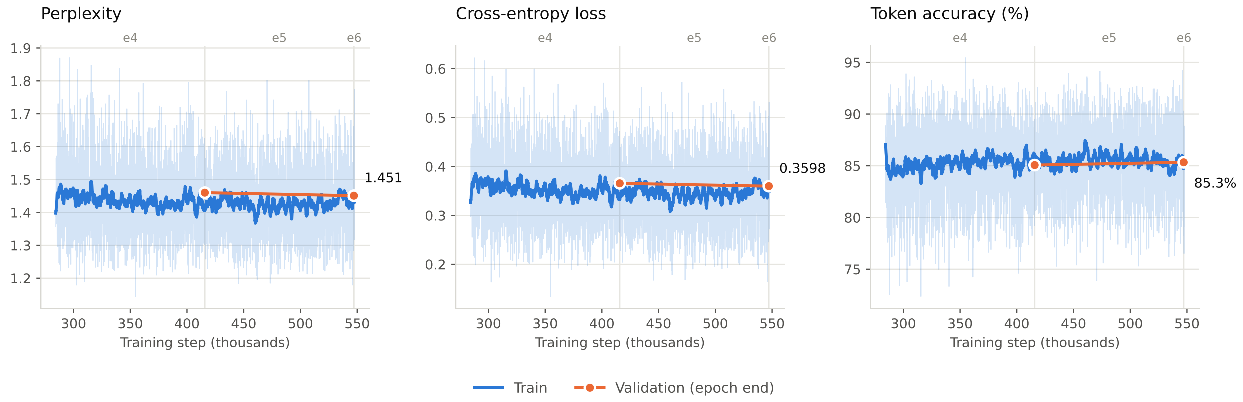


Figure 2: Continued pre-training on the whole, unclustered dataset, resumed from epoch 3.

Training used nucleotide-level tokenization with an 8-token vocabulary, including special tokens. The model was trained on 4 Nvidia A100 GPUs with bf16-mixed precision, using a per-device batch size of 16 and gradient accumulation over 4 steps, corresponding to an effective global batch size of 256 sequences per optimizer step. Optimization used AdamW with learning rate $5.0 \times 10^{-5}$, weight decay $0.01$, and $\beta = (0.9, 0.999)$. The learning rate schedule consisted of a 2000-step linear warmup followed by cosine decay to a minimum learning rate of $1.0 \times 10^{-5}$. Gradient clipping with maximum norm 1.0 was applied throughout training.

Figures 1 and 2 report perplexity, cross-entropy loss and token accuracy for the two pre-training stages. Both show stable optimization: the loss decreases smoothly after warmup with no divergence or loss spikes and the train-validation gap stays small and roughly constant, indicating convergence without overfitting. On the clustered dataset, validation perplexity falls from $1.72$ to $1.68$ and token accuracy reaches $77.8\%$ over four epochs. Continued training on the whole dataset, resumed from the final clustered checkpoint, reaches $1.45$ and $85.3\%$. The two are not directly comparable: the unclustered validation set retains the sequence redundancy that clustering removes, so homologous sequences make it easier to predict, and the lower perplexity reflects the evaluation set as much as further training.

##### A.3 Evaluation Metrics For Generated Sequences

**Normalized minimum free energy.** For each sequence $s$, ViennaRNA [5] is used to predict the secondary structure with minimum free energy, denoted by $\mathrm{MFE}(s)$. Because absolute free energy depends on sequence length, direct comparison across sequences of different lengths can be misleading. We therefore report the normalized minimum free energy

$$\mathrm{AMFE}(s) = \frac{\mathrm{MFE}(s)}{|s|},$$

where $|s|$ is the sequence length. This normalization allows thermodynamic stability to be compared on a per-nucleotide basis across datasets with different length distributions. More negative $\mathrm{AMFE}(s)$ values indicate more stable predicted folds, whereas values closer to zero indicate less stable structures.

**Novelty metric.** Novelty is evaluated by searching each generated sequence against the reference set with MMseqs2 `easy-search`. We use nucleotide-against-nucleotide search (`-search-type 3`) with sensitivity `-s 7.5`, retain up to 10,000 candidates after prefiltering (`-max-seqs 10000`), require a minimum sequence identity of 0.8 (`-min-seq-id 0.8`), and retain only hits with E-value at most $10^{-5}$ (`-e 1e-5`). We additionally use `-cov-mode 5` with `-c 0.9`, which enforces a strict compatibility constraint on sequence lengths by requiring the shorter sequence to span at least 90% of the length of the longer one. For each generated sequence $g$, we retain the best-scoring reported hit in the reference set. Let $\mathrm{alnlen}(g)$ denote the alignment length between $g$ and its best hit, let $|g|$ denote the length of the generated query sequence, let $\mathrm{pident}(g)$ denote the percentage of identical nucleotides in this alignment, and let $E(g)$ denote the E-value of the best hit for $g$. Smaller values of $E(g)$ indicate more statistically significant similarity to the reference set. We define the query coverage as

$$q_{\mathrm{cov}}(g) = \frac{\mathrm{alnlen}(g)}{|g|}.$$

Each generated sequence is then assigned to one of three novelty classes:

$$\mathrm{NoveltyClass}(g) = \begin{cases} \textit{known}, & E(g) \leq 10^{-10},\ q_{\mathrm{cov}}(g) \geq 0.7, \\ \textit{partially novel}, & g \text{ has a reported MMseqs2 hit, but is not } \textit{known}, \\ \textit{novel}, & g \text{ has no reported MMseqs2 hit.} \end{cases}$$

Under this definition, *known* sequences are high-confidence near-reference matches. *Partially novel* sequences retain detectable homology to the reference set, but do not satisfy the stricter criteria for the *known* class. Finally, *novel* sequences have no MMseqs2 hit satisfying the search criteria above and therefore occupy a more distant region of sequence space.

**Diversity metrics.** Diversity is evaluated by clustering sequences with MMseqs2 `easy-cluster`. In our implementation, clustering is performed with a minimum sequence identity threshold of 0.9 (`-min-seq-id 0.9`) and a coverage threshold of 0.8 (`-c 0.8`). MMseqs2 clustering was run under the default coverage mode (`-cov-mode 0`), under which the coverage threshold must be satisfied for both the query and target sequence. Let $N$ denote the total number of sequences, let $C$ denote the number of MMseqs2 clusters, let $N_{\mathrm{unique}}$ denote the number of unique sequences, and let $N_{\mathrm{singleton}}$ denote the number of clusters containing exactly one sequence. The reported diversity metrics are

$$\text{Diversity Ratio} = \frac{C}{N},$$

$$\text{Duplicate Rate} = \frac{N - N_{\mathrm{unique}}}{N} \times 100,$$

$$\text{Singleton Sequence Rate} = \frac{N_{\mathrm{singleton}}}{N} \times 100.$$

Here, the diversity ratio and singleton sequence rate are derived from the MMseqs2 clustering output, whereas the duplicate rate is computed separately from exact sequence identity counts. A higher diversity ratio and singleton rate indicate broader coverage of sequence space, while a higher duplicate rate indicates stronger redundancy.

**Levenshtein distance metric.** For aptamer experiments, the minimum normalized sequence Levenshtein distance for sequence $s_i$ is defined as

$$d_{\min}^{\text{seq}}(s_i) = \min_{j \neq i} \frac{\text{Lev}(s_i, s_j)}{\max(|s_i|, |s_j|)},$$

where $\text{Lev}(s_i, s_j)$ is the Levenshtein edit distance between sequences $s_i$ and $s_j$. Smaller values indicate that a sequence has a very close neighbor within the same generated set.

##### A.4 BEACON Tasks Hyperparameters

| Task | Head type | Dataset split (tr/val/te) | LR | Best epoch |
|---|---|---|---|---|
| Secondary structure | Nucleotide-Nucleotide Relation Prediction | 10814/1300/1305 | 1e-5 | 100.00 |
| Contact map | Nucleotide-Nucleotide Relation Prediction | 188/23/80+19+6 | 1e-5 | 100.00 |
| Distance map | Nucleotide-Nucleotide Relation Prediction | 188/23/80+19+6 | 1e-5 | 100.00 |
| Structural score imp. | Sequence Level (masked position regression) | 14049/1756/3095 | 1e-5 | 1.82 |
| SpliceAI | Nucleotide Level Prediction | 144628/18078/16505 | 1e-5 | 2.39 |
| Isoform | Sequence Level Prediction | 145463/33170/49755 | 1e-4 | 6.78 |
| NcRNA family | Sequence Level Prediction | 5670/650/2600 | 1e-4 | 13.54 |
| Modification | Sequence Level Prediction | 304661/3599/1200 | 1e-5 | 2.98 |
| Mean ribosome loading | Sequence Level Prediction | 76319/7600/7600 | 1e-4 | 10.31 |
| Degradation | Nucleotide Level Prediction | 1947/243/3005+3005 | 1e-5 | 9.84 |
| Programmable RNA switches | Sequence Level Prediction | 73227/9153/9154 | 1e-4 | 4.76 |
| CRISPR on-target | Sequence Level Prediction | 1453/207/416 | 1e-4 | 15.38 |
| CRISPR off-target | Sequence Level Prediction | 14223/2032/4064 | 1e-4 | 14.62 |

**Head definitions.**
We did not implement task-specific prediction heads ourselves. Instead, we adopted the BEACON benchmark evaluation pipeline and provided B-RANKO embeddings in the required input format for each task. The downstream heads therefore follow the standard BEACON setup.
*Sequence Level Prediction*: For sequence-level tasks, we use covariance pooling over nucleotide representations to obtain a sequence-level embedding. This representation is then processed through an MLP layer to derive the sequence-level predictions.
*Nucleotide Level Prediction*: For tasks requiring resolution at the nucleotide level, individual representations for each nucleotide are processed through an MLP to generate nucleotide-level predictions.
*Nucleotide-Nucleotide Relation Prediction*: To analyze relationships between nucleotides, we compute a self outer product of the nucleotide representations to form a matrix containing pairwise interactions between nucleotides. This matrix is then passed through a simple ResNet to get the final output.

#### B Aptamer Design Additional Datasets

Table 1: Summary of the AptaDiff train/validation splits. Sequence lengths are reported as mean ± standard deviation.

| Dataset | Train | Val | Train length ($\mu \pm \sigma$) | Val length ($\mu \pm \sigma$) |
|---|---|---|---|---|
| A (IGFBP3) | 12423 | 1380 | 36.0 ± 0.0 | 36.0 ± 0.0 |
| B (PTK7) | 10392 | 1155 | 36.0 ± 0.0 | 36.0 ± 0.0 |
| C | 58979 | 6553 | 84.9 ± 11.9 | 84.8 ± 11.8 |
| D | 76653 | 8517 | 87.4 ± 20.9 | 87.3 ± 20.9 |

#### B.1 Dataset B

Dataset B [6] was generated by the authors using an in-house HT-SELEX experiment targeting inactive tyrosine-protein kinase 7 (PTK7), with aptamer sequences consisting of a 36-nt variable region flanked by fixed primer regions. For dataset B (PTK7), B-RANKO was fine-tuned from the pretrained bidirectional checkpoint using a learning rate of $1 \times 10^{-5}$ for 3 epochs.

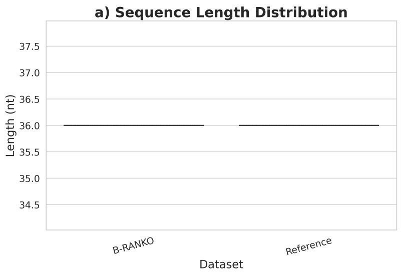


(a) Length distribution.

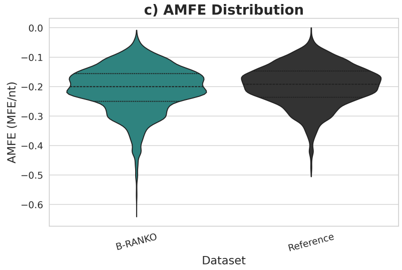


(b) MFE distribution.

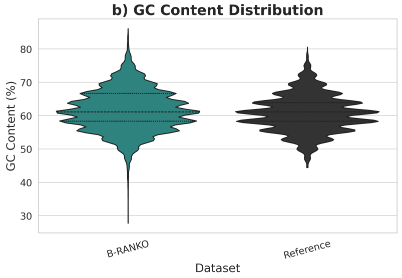


(c) GC content distribution.

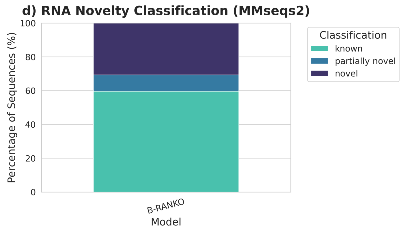


(d) Novelty classification.

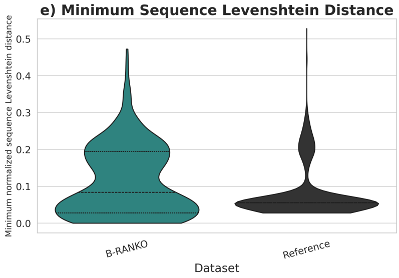


(e) Minimum sequence Levenshtein distance.

Figure 3: Evaluation of B-RANKO on AptaDiff dataset B. The panels summarize the distributions of sequence length, minimum free energy (MFE), and GC content, together with novelty and aptamer-specific diversity metrics based on minimum sequence and secondary-structure distances.

Table 2: MMseqs2 clustering-based diversity statistics for generated sequences on AptaDiff dataset B. Higher diversity ratio indicates broader coverage of sequence space, while a lower exact duplicate rate indicates lower redundancy.

| Model | Clusters | Diversity Ratio | Exact Duplicate Rate |
|---|---|---|---|
| B-RANKO | 4182 | 0.418 | 23.6% |
| Reference | 1436 | 0.144 | 0.0% |

#### B.2 Dataset C

Dataset C [6] is a public HT-SELEX dataset from DDBJ targeting human transglutaminase 2 (TGM2), with aptamer sequences containing a 30-nt variable region between fixed primer regions. For dataset C, B-RANKO was fine-tuned from the pretrained bidirectional checkpoint with a learning rate of $1 \times 10^{-5}$ for 3 epochs.

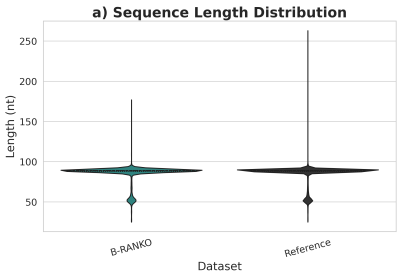


(a) Length distribution.

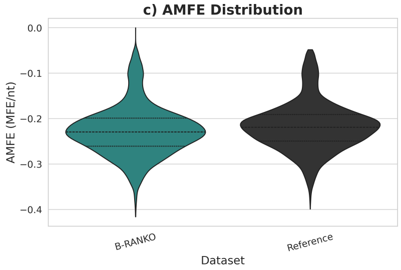


(b) MFE distribution.

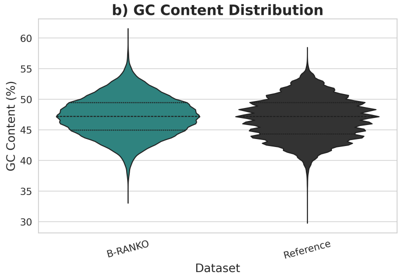


(c) GC content distribution.

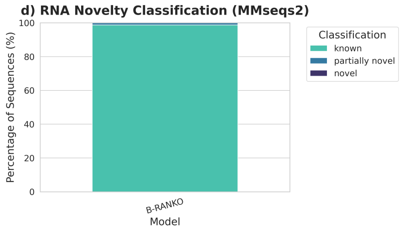


(d) Novelty classification.

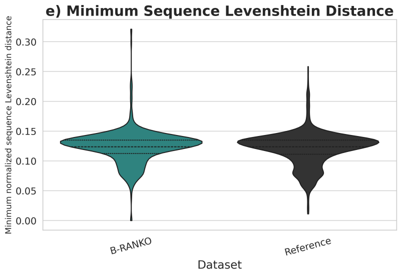


(e) Minimum sequence Levenshtein distance.

Figure 4: Evaluation of B-RANKO on AptaDiff dataset C. The panels summarize the distributions of sequence length, minimum free energy (MFE), and GC content, together with novelty and aptamer-specific diversity metrics based on minimum sequence and secondary-structure distances.

Table 3: MMseqs2 clustering-based diversity statistics for generated sequences on AptaDiff dataset C. Higher diversity ratio indicates broader coverage of sequence space, while a lower exact duplicate rate indicates lower redundancy.

| Model | Clusters | Diversity Ratio | Exact Duplicate Rate |
|---|---|---|---|
| B-RANKO | 9166 | 0.917 | 0.1% |
| Reference | 8798 | 0.880 | 0.0% |

#### B.3 Dataset D

Dataset D [6] is a public HT-SELEX dataset from DDBJ targeting human integrin $\alpha V \beta 3$, with aptamer sequences containing a 40-nt variable region flanked by fixed primer regions. For dataset D, B-RANKO was fine-tuned from the pretrained bidirectional checkpoint using a learning rate of $1 \times 10^{-5}$ for 3 epochs.

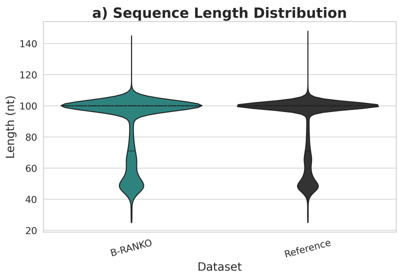


(a) Length distribution.

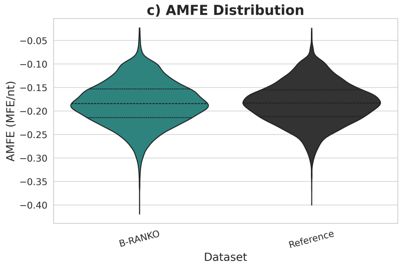


(b) MFE distribution.

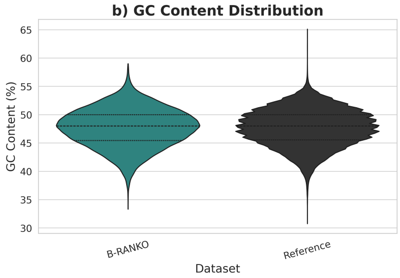


(c) GC content distribution.

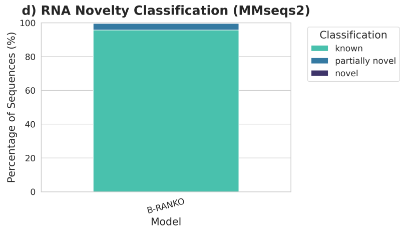


(d) Novelty classification.

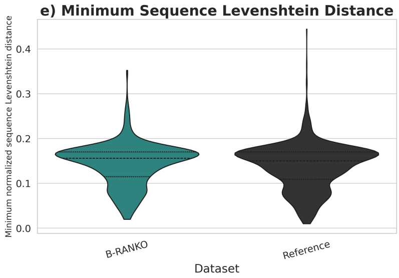


(e) Minimum sequence Levenshtein distance.

Figure 5: Evaluation of B-RANKO on AptaDiff dataset D. The panels summarize the distributions of sequence length, minimum free energy (MFE), and GC content, together with novelty and aptamer-specific diversity metrics based on minimum sequence and secondary-structure distances.

Table 4: MMseqs2 clustering-based diversity statistics for generated sequences on AptaDiff dataset D. Higher diversity ratio indicates broader coverage of sequence space, while a lower exact duplicate rate indicates lower redundancy.

| Model | Clusters | Diversity Ratio | Exact Duplicate Rate |
|---|---|---|---|
| B-RANKO | 9072 | 0.907 | 0.1% |
| Reference | 8018 | 0.802 | 0.0% |

### C Ablation Studies

#### C.1 Additional BEACON baselines

Table 5: Additional published BEACON reference results.

| Task | SSP | CMP | DMP | SSI | SPL | APA | NcRNA | Modif | MRL | VDP | PRS | CRI-On | CRI-Off |
|---|---|---|---|---|---|---|---|---|---|---|---|---|---|
| Metric | F1 (%) | P@L (%) | $R^2$ (%) | $R^2$ (%) | ACC@K (%) | $R^2$ (%) | ACC (%) | AUC (%) | $R^2$ (%) | MCRMSE↓ | $R^2$ (%) | SC (%) | SC (%) |
| *Literature SOTA* | | | | | | | | | | | | | |
| Method | UFold | RNACon | SS+Seq | StructImp | SpliceAI | APARENT | GCN | MultiRM | Optimus | NAttn | MLP-O | SSC | DeepCRI |
| SOTA | 65.40 | 66.00 | 68.75 | 37.20 | 32.18 | 50.82 | 85.73 | 84.00 | 78.00 | 0.263 | 55.67 | 44.10 | 12.60 |
| *Naive supervised models* | | | | | | | | | | | | | |
| CNN | 49.95 | 43.89 | 27.76 | 34.36 | 8.43 | 50.93 | 88.62 | 70.87 | 74.13 | 0.361 | 45.40 | 29.69 | 11.40 |
| ResNet | 57.26 | 59.59 | 30.26 | 37.74 | 21.15 | 56.45 | 88.32 | 71.03 | 74.34 | 0.349 | 55.21 | 28.55 | 11.50 |
| LSTM | 58.61 | 40.41 | 44.77 | 35.44 | 36.66 | 67.03 | 88.78 | 94.83 | 83.94 | 0.329 | 55.45 | 26.83 | 8.60 |
| *Pretrained BEACON baseline* | | | | | | | | | | | | | |
| BEACON-B | 64.18 | 60.81 | 56.28 | 38.78 | 37.43 | 70.59 | 94.63 | 94.74 | 72.29 | 0.320 | 54.67 | 26.01 | 4.42 |
| BEACON-B512 | 58.75 | 61.20 | 56.82 | 39.13 | 37.24 | 72.00 | 94.99 | 94.92 | 72.35 | 0.320 | 55.20 | 28.17 | 3.82 |

All values are taken from the BEACON paper [7] and were not re-evaluated by us. As these are single reported values without seed variation, small differences relative to our $\text{mean}_{(\text{sd})}$ results should not be interpreted as meaningful. The literature SOTA row contains task-specific supervised models trained and tuned for individual tasks, and therefore serves as a reference rather than a direct comparison with general-purpose foundation models.

#### C.2 Pooling: mean versus covariance

Sequence-level prediction tasks require pooling token representations into a single vector. We compare mean pooling over non-special tokens with the covariance pooler described in Section 2.5, while keeping the backbone, task head, number of training epochs, and random seeds fixed.

Table 6: Mean versus covariance pooling on the seven pooled BEACON tasks. Values are $\text{mean}_{(\text{sd})}$ over seeds 42, 43 and 44. $\Delta$ is covariance minus mean.

| Task | Mean pooling | Covariance pooling | $\Delta$ |
|---|---|---|---|
| APA [$R^2$ %] | $69.52_{(0.58)}$ | $69.11_{(1.00)}$ | $-0.41$ |
| NcRNA [ACC %] | $95.12_{(0.12)}$ | $96.72_{(0.59)}$ | $+1.60$ |
| Modif [AUC %] | $95.04_{(0.03)}$ | $95.08_{(0.11)}$ | $+0.03$ |
| MRL [$R^2$ %] | $84.44_{(0.11)}$ | $85.04_{(0.25)}$ | $+0.60$ |
| PRS [$R^2$ %] | $57.13_{(0.18)}$ | $57.67_{(0.16)}$ | $+0.54$ |
| CRI-On [SC %] | $23.11_{(3.51)}$ | $26.27_{(6.48)}$ | $+3.16$ |
| CRI-Off [SC %] | $8.85_{(2.71)}$ | $10.77_{(1.59)}$ | $+1.92$ |

Mean and covariance pooling perform similarly across the evaluated tasks, with covariance pooling showing a marginal overall advantage. While the gains vary across tasks, covariance pooling improves most of the evaluated tasks and achieves larger improvements on ncRNA, MRL, PRS, and the two CRISPR tasks. We therefore use covariance pooling in our experiments.

#### C.3 Extraction attention: full versus inward

B-RANKO is pretrained using bidirectional autoregressive attention, while downstream representations are extracted using full bidirectional attention. Since this attention pattern is not used during pretraining, we re-ran all 13 tasks using the training-time mask for representation extraction, with each sequence's centre, $\lfloor T/2 \rfloor$, used as the merge index to match the inference default.

Table 7: Full bidirectional attention versus the training time Bidirectional autoregressive attention, using mean pooling. Results use seeds 42,43,44 and $\text{mean}_{(sd)}$ are reported. $\Delta$ denotes bidirectional autoregressive minus full bidirectional attention.

| Task | Full bidirectional attention | Bidirectional autoregressive attention | $\Delta$ |
|---|---|---|---|
| SSP [F1 %] | $49.12_{(0.26)}$ | $46.08_{(2.70)}$ | $-3.04$ |
| CMP [P@L %] | $44.66_{(0.98)}$ | $45.98_{(0.56)}$ | $+1.32$ |
| DMP [$R^2$ %] | $57.78_{(0.52)}$ | $57.34_{(0.16)}$ | $-0.43$ |
| SSI [$R^2$ %] | $39.69_{(0.36)}$ | $21.51_{(0.12)}$ | $-18.18$ |
| SpliceAI [ACC@K %] | $39.38_{(1.09)}$ | $18.49_{(0.74)}$ | $-20.89$ |
| APA [$R^2$ %] | $69.52_{(0.58)}$ | $61.23_{(2.42)}$ | $-8.29$ |
| NcRNA [ACC %] | $95.12_{(0.12)}$ | $94.14_{(0.32)}$ | $-0.97$ |
| Modif [AUC %] | $95.04_{(0.03)}$ | $73.39_{(0.46)}$ | $-21.66$ |
| MRL [$R^2$ %] | $84.44_{(0.11)}$ | $75.86_{(0.28)}$ | $-8.58$ |
| VDP [MCRMSE $\downarrow$] | $0.264_{(0.003)}$ | $0.347_{(<0.001)}$ | $+0.084$ |
| PRS [$R^2$ %] | $57.13_{(0.18)}$ | $57.20_{(0.24)}$ | $+0.06$ |
| CRI-On [SC %] | $23.11_{(3.51)}$ | $36.05_{(2.67)}$ | $+12.93$ |
| CRI-Off [SC %] | $8.85_{(2.71)}$ | $9.73_{(0.83)}$ | $+0.89$ |

Full bidirectional attention generally performs better than bidirectional autoregressive attention for representation extraction. The largest differences appear on SSI, SpliceAI, Modif, APA, and MRL, while the two approaches perform similarly on CMP, DMP, NcRNA, and PRS. Although bidirectional autoregressive attention improves both CRISPR tasks, the overall results favor full bidirectional attention. We therefore use full bidirectional attention for downstream representation extraction, while retaining bidirectional autoregressive attention for sequence generation.

##### C.4 Configuration of the matched-scale comparison

Table 8 lists the configuration of the two models compared in Section 3.4. The backbones are parameter-identical. The only difference in weights is the second output head of the bidirectional model, $480 \times 8 = 3{,}840$ parameters.

Table 8: Configuration of the controlled mask ablation.

| | Bidirectional | Causal |
|---|---|---|
| Tokenizer | single-nucleotide (8) | single-nucleotide (8) |
| Width / blocks / heads | 480 / 12 / 20 | 480 / 12 / 20 |
| Trunk parameters | 33.2373 M | 33.2373 M |
| Trainable parameters | 33.2498 M | 33.2460 M |
| Pretraining corpus | Rfam representatives | Rfam representatives |
| Epochs / optimizer steps | 10 / 101,760 | 10 / 101,760 |
| Effective batch | 256 | 256 |
| Optimizer, schedule, precision | identical | identical |
| Downstream pooling | mean | mean |

#### References

[1] Marena I. Trinidad and Jamie H. D. Cate. GARNET database, 2024. URL `https://doi.org/10.5281/zenodo.14642546`. Data retrieved from Zenodo repository.

[2] Ioanna Kalvari, Eric P. Nawrocki, Nancy Ontiveros-Palacios, Joanna Argasinska, Kevin Lamkiewicz, Manja Marz, Sam Griffiths-Jones, Claire Toffano-Nioche, Daniel Gautheret, Zasha Weinberg, Elena Rivas, Sean R. Eddy, Robert D. Finn, Alex Bateman, and Anton I. Petrov. Rfam 14: expanded coverage of metagenomic, viral and microRNA families. *Nucleic Acids Research*, 49(D1):D192–D200, 2021. doi: 10.1093/nar/gkaa1047. URL `https://doi.org/10.1093/nar/gkaa1047`.

[3] The RNAcentral Consortium. RNAcentral 2021: secondary structure integration, improved sequence search and new member databases. *Nucleic Acids Research*, 49(D1):D212–D220, 2021. doi: 10.1093/nar/gkaa921. URL `https://doi.org/10.1093/nar/gkaa921`.

[4] Martin Steinegger and Johannes Söding. MMseqs2 enables sensitive protein sequence searching for the analysis of massive data sets. *Nature Biotechnology*, 35(11):1026–1028, 2017. doi: 10.1038/nbt.3988. URL `https://doi.org/10.1038/nbt.3988`.

[5] Ronny Lorenz, Stephan H Bernhart, Christian Höner zu Siederdissen, Hakim Tafer, Christoph Flamm, Peter F Stadler, and Ivo L Hofacker. Viennarna package 2.0. *Algorithms for Molecular Biology*, 6(1):26, 2011. doi: 10.1186/1748-7188-6-26.

[6] Z. Wang et al. Aptadiff: de novo design and optimization of aptamers based on diffusion models. *Briefings in Bioinformatics*, 25(6):bbae517, 2024. doi: 10.1093/bib/bbae517. URL `https://doi.org/10.1093/bib/bbae517`.

[7] Yuchen Ren, Zhiyuan Chen, Lifeng Qiao, Hongtai Jing, Yuchen Cai, Sheng Xu, Peng Ye, Xinzhu Ma, Siqi Sun, Hongliang Yan, Dong Yuan, Wanli Ouyang, and Xihui Liu. Beacon: Benchmark for comprehensive rna tasks and language models. *arXiv preprint arXiv:2406.10391*, 2024. URL `https://arxiv.org/abs/2406.10391`.
